# Decoding the Influence of Central LEAP2 on Hedonic Food Intake and its association with Dopaminergic Reward Pathways

**DOI:** 10.1101/2023.08.29.555294

**Authors:** Maximilian Tufvesson-Alm, Qian Zhang, Cajsa Aranäs, Sebastian Blid Sköldheden, Christian E Edvardsson, Elisabet Jerlhag

## Abstract

The gut-brain peptide ghrelin and its receptor (GHSR) are established as a regulator of hunger and reward-processing. However, the recently recognized GHSR inverse agonist, liver-expressed antimicrobial peptide 2 (LEAP2), is less characterized. Given the role of GHSR in many central processes, and in particular reward, understanding the central effects of LEAP2 is of high interest to understand reward-related behaviors and disorders, including hedonic feeding in eating disorders. The present study aimed to elucidate LEAP2s central effect on reward-related behaviors through hedonic feeding and its mechanism. LEAP2 was administrated centrally in male mice and effectively reduced hedonic feeding but had no or little effect on homeostatic chow intake when a more palatable option was available. Strikingly, the effect on hedonic feeding was correlated to the preference of the palatable food option, where peanut butter showed the highest preference and the greatest reduction by LEAP2. Further, LEAP2 reduced the rewarding memory of high-preference foods, as well as attenuated the accumbal dopamine release associated with peanut butter exposure and eating. Interestingly, LEAP2 was widely expressed in the brain, and in particular in reward-related brain areas such as the laterodorsal tegmental area (LDTg). The expression in this area was also markedly altered when given free access to peanut butter. Accordingly, infusion of LEAP2 into the LDTg was sufficient to attenuate acute peanut butter eating. Taken together, the present results show that central LEAP2 has a profound effect on central dopaminergic reward signaling and affects several aspects of hedonic eating. The present study highlights LEAP2s effect on reward, which may have application not only for hedonic feeding, but for other reward-related psychiatric disorders as well.

## Introduction

Liver-expressed antimicrobial peptide 2 (LEAP2) was recently recognized as the endogenous antagonist to ghrelin, working as an inverse agonist on growth hormone secretagogue receptor (GHSR)^1, 2^. Ghrelin and GHSR have a crucial role in feeding, and an increasingly recognized role in reward processing^3, 4^. In this regard ghrelin increases, whereas synthetic GHSR antagonists attenuates reward-related behavior^5–11^. In particular, GHSR modulates the cholinergic-dopaminergic reward pathway. Thus, activation of cholinergic afferents from laterodorsal tegmental area (LDTg) to ventral tegmental area (VTA), and VTA neurons directly, result in dopamine release in the nucleus accumbens (NAc) shell^8, 9, 12, 13^. As such, understanding the central effects of LEAP2, and thus the whole ghrelinergic system, is of high interest for many reward-related behaviors^14^. This includes food intake in eating disorders^15, 16^ and psychiatric disorders with a disrupted reward processing^17–19^. However, reports regarding LEAP2s effect on food intake have been inconsistent, where some studies have shown that LEAP2 reduces regular chow intake^1^, and some reporting it does not affect chow intake, but suppresses ghrelin-induced food intake^20, 21^. Interestingly, there appears to be a discrepancy between central and systemic administration, where central, but not systemic, LEAP2 specifically reduces high-fat diet intake^20^, indicating that LEAP2 regulate hedonic food intake through central processes. Although LEAP2 is mainly produced by the small intestine and liver^22^, LEAP2 has been shown to be expressed in the mouse brain, including hypothalamus, midbrain and hippocampus^21^, and can be detected in human cerebrospinal fluid (CSF)^23^. This raises the possibility that LEAP2 has a function within the brain to influence food intake through central reward processes via the GHSR. Indeed, LEAP2 has been shown to alter dopaminergic signaling through the heterodimerization of GHSR-D_2_ receptor complex *in vitro*, further suggesting its involvement in modulating the reward system^24^. Additionally, a recent study showed that central, but not systemic administration of LEAP2 reduces binge-like alcohol intake in mice, further strengthening LEAP2s central role^25^. However, information regarding LEAP2s function, and in particular its effect on central reward circuitry and the behavioral outcome, is sparse.

Taken together, it appears credible that LEAP2 would have considerable central effects, in particular on the dopaminergic reward system, reward-related behaviors and hedonic feeding. In the present study, we sought to clarify LEAP2s central effect on hedonic food intake and investigate its influence over the central reward system and reward-driven behaviors.

## Material and methods

### Animals

Adult male NMRI mice (25–30 g body weight at arrival; Charles River; Sulzfeld, Germany) were used. The mice were group-housed and habituated to the animal facility one week before exposure to the hedonic food: peanut butter (Crunchy, Green choice, Sweden; 5.9 kcal/g), Nutella® (Ferrero, Pino Torinese, Italy; 5.5 kcal/g) or chocolate (Milk Chocolate, Marabou, Upplands Väsby, Sweden; 5.5 kcal/g). Mice were exposed group housed for two days and then separated to allow for individual measuring. In addition to hedonic foods, all animals had free access to standard chow (Teklad Rodent Diet; Envigo, Madison, WI, USA; 3 kcal/g) and water. They were kept in a room with a 12-h light dark cycle (lights on at 7 a.m.) with a temperature of 20 °C and humidity of 50%. Mice were handled on three occasions before experiment and always habituated to the experimental room for one hour. Following experiment conclusion, mice were sacrificed, and the brains collected. Only animals with correct placement for injection and probe were included (Supplementary Figure 1). All experiments were conducted in accordance with guidelines from the Swedish Ethical Committee on Animal Research in Gothenburg (ethical permits 1457/18 and 3348/20) and every effort was made to maximize the animal’s well-being.

### Drugs and administration

LEAP2 (LEAP-2 (38-77) (Human) / LEAP-2 (37-76) (Mouse), 075-40, Phoenix Pharmaceuticals Inc., USA) was dissolved in vehicle (Ringer solution; NaCl 140 mM, CaCl_2_1.2 mM, KCl 3 mM and MgCl_2_ 1 mM (Merck KGaA Darmstadt, Germany)) to a dilution of 5.5μg/μl. Two days prior to start of any experiment, animals underwent surgery to implant guide cannula for intracerebroventricular (i.c.v.) injection or local injection into the LDTg (Supplementary Figure 1A and C), as previously described^26, 27^. One hour prior to central infusion start a dummy injector targeting the infusion area was inserted into the guide cannula and then retracted to remove clotted blood and hamper spreading depression. Fifteen minutes before testing, 1μl of the solution or vehicle was slowly injected over 1 minute in the third ventricle or 0.5μl bilaterally into the LDTg and left in place for another minute before being retracted to allow for complete diffusion of the drug. The dose (5.5μl/mouse) was selected based on dose-response studies where it was the highest tested dose and did not affect the normal state of the animal measured by locomotor activity (Supplementary Figure 2).

### Food intake experiments

The potential of central LEAP2 to reduce the intake of hedonic foods and chow was tested in a series of experiments where LEAP2 or vehicle was infused into the third ventricle. Hedonic food consumption, chow consumption, total consumption and preference, calculated as percentage of hedonic food intake compared to total food intake in weight, was measured 1h, 2h, 4h and 24h following treatment. To test the effect of LEAP2 in LDTg, hedonic food was only available for 2 hours every day to simulate acute rewarding effect and food intake was only measured for those hours following bilateral administration of LEAP2 or vehicle. The treatments were randomized between high- and low-consumers of hedonic foods, as measured in the days before experimental start. After the first treatment (vehicle or LEAP2) the opposite treatment was infused the following day, to allow for paired comparisons. Importantly, the intake of vehicle treated mice was similar independent of treatment day.

### Conditioned place preference

The effect of central LEAP2 on reward-related behaviors is currently unexplored, but considering its effect on food intake, it appears likely that food-reward is affected. Here, reward-dependent memory of hedonic foods was measured in the memory conditioned place preference (mCPP) paradigm as previously described^28^. During preconditioning (day 1), mice allowed to freely explore both compartments for 20 minutes. The least preferred compartment was then paired with hedonic foods in a biased approach and the other chamber was empty for two conditioning sessions per day over four days (day 2-5). Animals were then administered vehicle or LEAP2 i.c.v. on day 6 and mCPP was calculated as the difference of the total time spent in the food-paired compartment during the post-conditioning and pre-conditioning sessions as a percent of the total time.

### Microdialysis and dopamine analysis

The central processes by which LEAP2 affects hedonic food intake and reward is currently unknown. Here, LEAP2s effect on dopaminergic reward signaling in association with food was tested using microdialysis. Two days prior to microdialysis experiment, mice pre-exposed to peanut butter underwent surgery as previously described^29^. A guide cannula targeting third ventricle and a probe (20 kDa cut off with a 1 mm exposed length, HOSPAL, Gambro, Lund, Sweden) aiming at NAcS was inserted (Supplementary Figure 1B).

Dopaminergic signaling was measured using microdialysis as previously described^29^. In short, the probe implanted in freely moving animals, pre-exposed to peanut butter, was connected to a pump and perfused with Ringer’s solution at a rate of 1.6μl/min and samples were collected every 20 minutes. After a two-hour wash-out period, a baseline was collected (−80 to 0 min) followed by administration of vehicle or LEAP2 (after 0 min sample collected) and the presentation of peanut butter enclosed under a metal wire mesh cup which allowed for visual and olfactory stimulation (after 20 min sample collected), or a pencil cup alone for control animals. After the 60 min sample was collected, for a subset of the animals, the pencil cup was removed, and the animals allowed to eat.

The dopamine content of the samples was quantified using high-performance liquid chromatography system with electrochemical detection as described previously, according to a modified protocol^29, 30^. The dopamine levels were calculated as a percentage of the mean of the three baseline values before treatment. Additionally, the peanut butter exposure response was calculated as the percent change following presentation (40 min) from the sample just before presentation (20 min). Further, the area under the curve for eating from just before (60 min) was calculated.

### qPCR

Although LEAP2 has been shown to be expressed in larger brain areas, including hypothalamus, hippocampus and midbrain, a detailed mapping of reward-related areas is lacking. Further, how central LEAP2 expression is affected by diet and reward is currently unknown. Therefore, LEAP2 expression was quantified in the brain of animals that had free access to peanut butter or normal diet as a control. After one week the animals were sacrificed and the brains, as well as the liver and duodenum in a few control mice, were immediately frozen on dry ice and stored in −80°C. While kept on ice, the brains were cut in 1mm thick slices using a mouse brain matrix and the following areas were punched out: prefrontal cortex (PFC), NAc, bed nucleus of the stria terminalis (BNST), hippocampus, amygdala, hypothalamus, VTA, dorsal raphe (DR) and LDTg. Areas were pooled in groups of three in order to ensure sufficient amount of RNA to reliably measure expression. Total RNA was extracted, purified and amplified as done before^29, 31, 32^. The expression of the *leap2* gene (ThermoFisher, Mm00461982) was normalized to the geometric mean of *beta-actin* (ThermoFisher, Mm01205647). The comparative CT method (ABI technical manual) was used to analyze the real-time PCR. In control animals, LEAP2 expression was firstly compared and normalized to hypothalamus expression and shown as fold change, while the Δ_CT_-values were used to determine any deviation from hypothalamus expression or comparing control to peanut butter-eating mice within each area.

### Statistics

For all statistical calculations GraphPad Prism® 9.5.1. (GraphPad Software, Inc. La Jolla, CA, USA) was used. Gaussian distribution was tested for using D’Agostino and Pearson normality test and non-parametric tests used where appropriate. All test where two-tailed and alfa was set to 0.05. For feeding experiments, group comparisons were made using repeated measures two-way ANOVA with Wilcoxon signed rank test post-hoc. Hedonic foods 24-hour preference differences were analyzed using Kruskal-Wallis test followed by Mann-Whitney U test post-hoc. Correlation was analyzed using Spearman correlation. In CPP experiments comparisons were made using unpaired t-test. For microdialysis, group differences were assessed using mixed-effect analysis with Dunnet’s multiple comparisons test post hoc and using one-way ANOVA followed by Šidák’s multiple comparisons test post hoc. Expression of LEAP2 in different areas was analyzed using one-way ANOVA followed by Šidák’s multiple comparisons test post hoc and mice with free access to peanut butter were compared to the same area in control mice using unpaired t-test.

## Results

### Central administration of LEAP2 reduces consumption of hedonic foods without affecting chow consumption

In the food experiment, two-way repeated-measures ANOVA revealed that mice consumed significantly less peanut butter (Figure 1A, n=9, Treatment effect: F_(1, 8)_=8.38, p=0.020, Time effect: F_(1.04, 8.32)_=3.28, p=0.11, Interaction: F_(1.48, 11.83)_=5.480, p=0.028), Nutella® (Figure 1D, n=10, Treatment effect: F_(1, 9)_=5.87, p=0.038, Time effect: F_(1.03, 9.29)_=6.23, p=0.033, Interaction: F_(1.04, 9.36)_= 0.022, P=0.89) and chocolate (Figure 1G, n=11, Treatment effect: F_(1, 10)_=8.03, p=0.018, Time effect: F_(1.03, 10.25)_=3.93, p=0.074, Interaction: F_(1.02, 10.20)_=1.68, p=0.22) after administration of LEAP2 i.c.v. compared to vehicle. This effect was evident at two hours for Nutella® (p=0.017), up to two hours for chocolate (p=0.016 and p=0.012 for one and two hours, respectively) and an effect evident up to four hours for peanut butter (p=0.016, p=0.012 and p=0.027 for one, two and four hours respectively).

**Figure 1.**
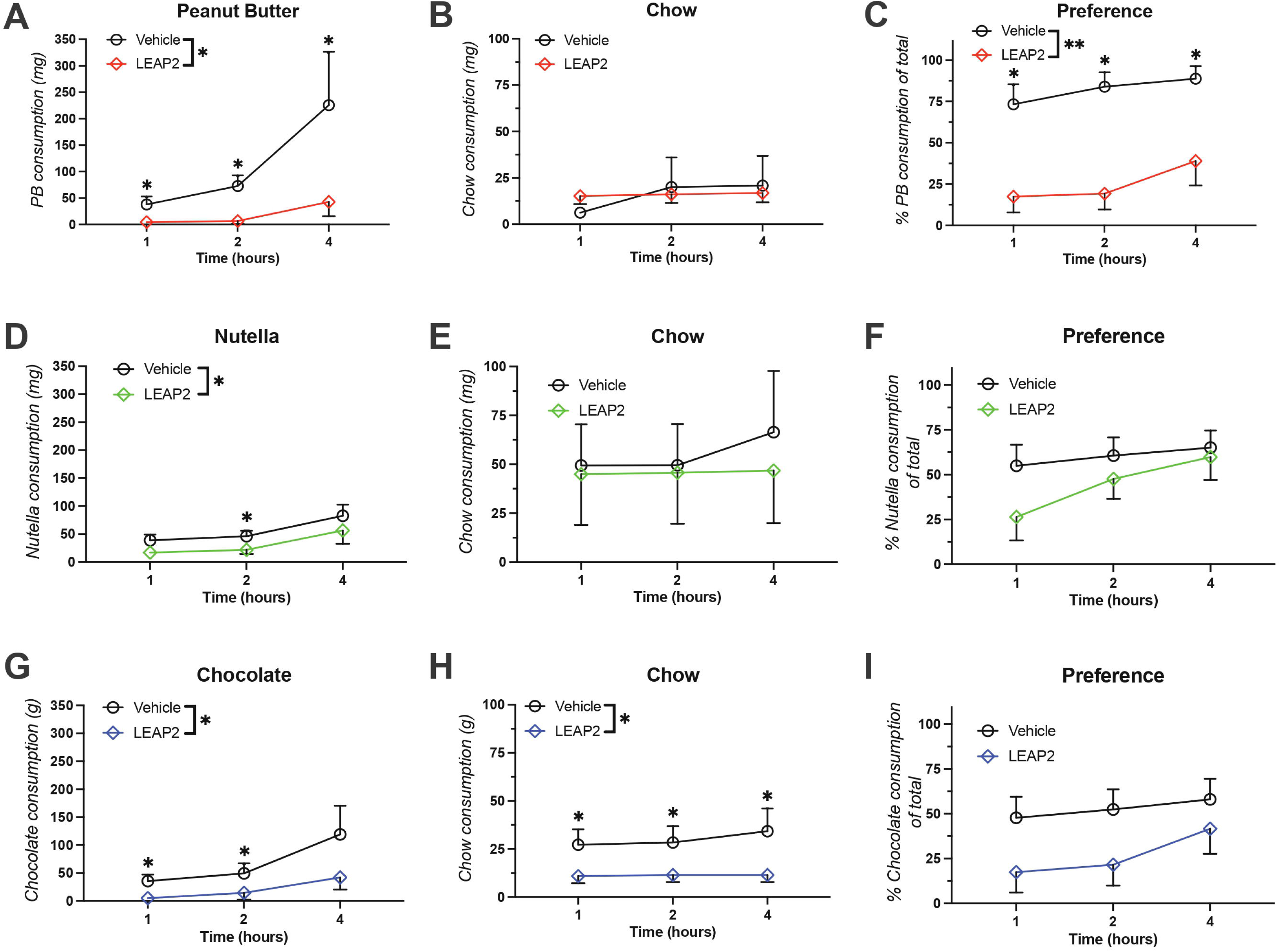
Central LEAP2 reduces hedonic feeding. The effect of centrally administered LEAP2 was assessed for three different hedonic foods, chow and the preference for the hedonic foods. For mice receiving peanut butter (A-C, n=9), LEAP2 reduced hedonic food intake and preference, but had no effect on chow intake. For Nutella® (D-F, n=10), a weaker effect was observed on hedonic food intake, but no effect on chow intake or on preference. Mice receiving chocolate (G-I, n=11) on the other hand showed reduction in both hedonic food and chow intake, but no difference in preference. Group comparisons were made using repeated measures two-way ANOVA with Wilcoxon signed rank test post-hoc. Data is shown as mean±SEM and was corrected for multiple comparison. p**<0.01, p*<0.05 vs. corresponding vehicle group.

No difference in chow intake was evident for peanut butter (Figure 1B, Treatment effect: F_(1, 8)_=0.0011, p=0.97, Time effect: F_(1.00, 8.03)_=1.20, p=0.305, Interaction: F_(1.01, 8.04)_=0.91, p=0.37) or Nutella®-fed mice (Figure 1E, Treatment effect, F_(1, 9)_=0.071, p=0.80, Time effect: F_(1.00, 9.01)_=1.35, p=0.27, Interaction: F_(1.00, 9.005)_=0.9081, p=0.37). However, in chocolate-fed animals LEAP2 treatment reduced chow intake (Figure 1H, Treatment effect: F_(1, 10)_=6.17, p=0.032, Time effect: F_(1.08, 10.83)_=2.01, p=0.18, Interaction: F_(1.08, 10.83)_=2.01, p=0.18, Interaction: F_(1.09, 10.91)_=1.58, p=0.24), which was evident up to four hours after administration (p=0.040 for all timepoints).

Further, a reduction in preference was only shown for peanut butter (Figure 1C, Treatment effect: F_(1, 8)_=23.25, p=0.0013, Time effect: F_(1.76, 14.07)_=7.512, p=0.0074, Interaction: F_(1.28, 10.24)_=0.78, p=0.43) and evident up to four hours (p=0.016, p=0.012 and p=0.027 for one, two and four hour timepoints, respectively), whereas preference for Nutella® (Figure 1F, Treatment effect: F_(1, 9)_=1.79, p=0.21, Time effect: F_(1.32, 11.90)_=6.09, p=0.023, Interaction: F _(1.18, 10.63)_=3.10, p=0.103) and chocolate (Figure 1I, Treatment: F_(1, 10)_=4.41, p=0.062, Time effect: F_(1.04, 10.37)_= 5.31, p=0.042, Interaction: F_(1.260, 12.56)_=1.60, p=0.24) was unaffected by LEAP2 administration.

### The effect of LEAP2 on hedonic food intake is dependent on preference

Given that the largest reduction in hedonic food intake was seen for peanut butter, which also had the highest preference, the association thereof was further investigated. During a 24-hour period of free access to hedonic foods and chow Kruskal-Wallis test revealed a significant group difference (p=0.0067) where peanut butter showed the highest preference (Figure 2A, 81.0±7.9%) and was significantly lower for Nutella® (63.1±3.3%, p=0.044) and chocolate (54.6±4.8%, p=0.0084). The reduction in hedonic food intake four hours after LEAP2 administration showed a significant positive correlation with 24-hour hedonic food preference (Figure 3B, p=0.0040, r=0.51).

**Figure 2.**
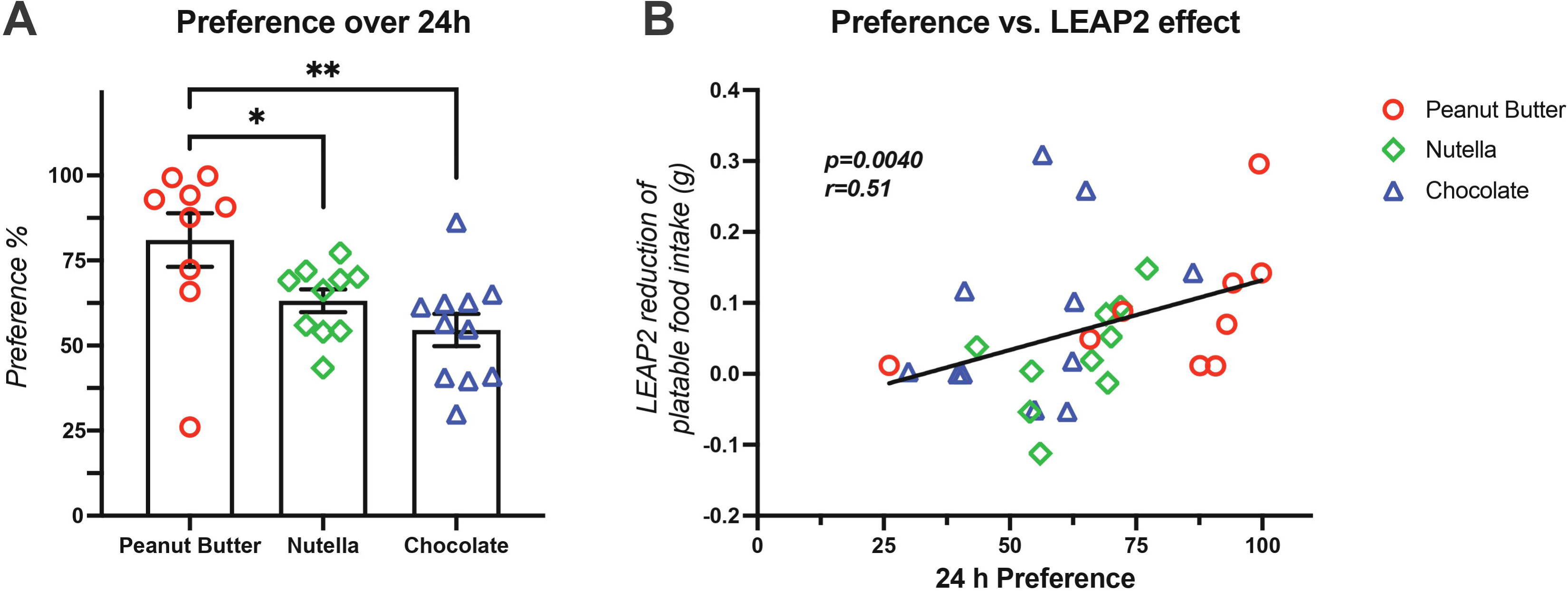
LEAP2s effect on hedonic foods are dependent on preference. The 24-hour preference for the hedonic foods and its correlation to the effect of LEAP2. Peanut butter (A, n=9) showed the highest preference followed by Nutella® (n=10) and chocolate (n=11). The reduction in hedonic food intake four hours after LEAP2 administration (B) was positively associated with the 24-hour preference. Hedonic foods 24-hour preference differences were analyzed using Kruskal-Wallis test followed by Mann-Whitney U test post-hoc. Correlation was analyzed using Spearman correlation. Data is shown as mean±SEM and was corrected for multiple comparison. p**<0.01, p*<0.05 vs. corresponding group.

**Figure 3.**
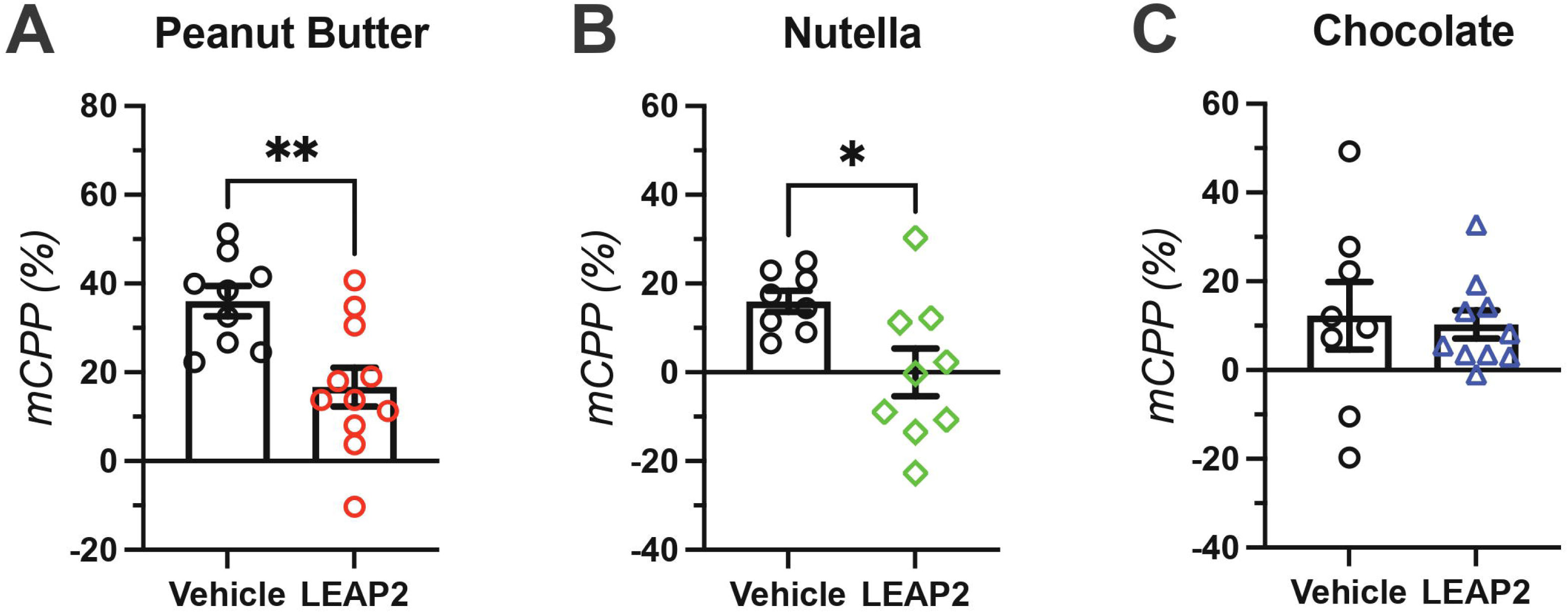
Central LEAP2 reduced rewarding memory of hedonic feeding. Effect of central LEAP2 on memory in the conditioned place preference. Central LEAP2 administration was able to attenuate the rewarding memory of peanut butter (A, n=9 and n=11 for vehicle and LEAP2, respectively) and Nutella® (B, n=8 and n=9 for vehicle and LEAP2, respectively), but not chocolate (C, n=8 and n=10, for vehicle and LEAP2, respectively). Comparisons were made using unpaired t-test. Data is shown as mean±SEM. p**<0.01, p*<0.05 vs. corresponding vehicle group.

### Central LEAP2 reduces the rewarding memory of hedonic food intake

Next, LEAP2s effect on reward-related memory retrieval associated hedonic food intake was evaluated using the mCPP paradigm where pre-exposed mice were only allowed to eat hedonic foods during training in the associated chamber. As expected, considering the preference and reduction of LEAP2 seen in the *ad libitum* food experiments, peanut butter caused the strongest CPP in vehicle-treated animals (Figure 3A, 36.0±3.4%, n=9), which was reduced by i.c.v. LEAP2 administration (16.7±4.4%, n=11, p=0.0035). LEAP2 was additionally able to fully attenuate CPP caused by Nutella® (Figure 3B, 0.0±5.4%, n=9) compared to vehicle (16.0±2.4%, n=8, p=0.020). However, LEAP2 administration had no effect on the CPP caused by chocolate (Figure 3C, 12.3±7.7%, n=8 and 10.3±3.2%, n=10 for vehicle and LEAP2 treatment, respectively, p=0.80).

### Central LEAP2 abolishes the accumbal dopamine release associated with peanut butter exposure and consumption

As peanut butter showed the highest amount of consumption, preference, and strongest rewarding memory, it was chosen for further investigation of LEAP2s effect on hedonic feeding. In order to explore LEAP2s effect on reward, microdialysis was used to measure the dopaminergic response to peanut butter exposure and eating. Here, a mixed-effects analysis revealed a significant group difference when exposed to peanut butter (Figure 4A, Treatment effect: F_(3, 25)_=6.17, p=0.0028, Time effect: F_(3.26, 70.18)_=13.20, p<0.0001, Interaction: F_(27, 194)_=2.40, p=0.0003). Further analysis revealed a significant group difference in the dopamine level response to peanut butter exposure compared to directly before (Figure 4B, one-way ANOVA, F_(3, 25)_=7.447, p=0.0010), where vehicle treated mice showed a significant increase of dopamine (50.7±10.4%, n=9) compared to LEAP2 treated (1.3±7.0%, n=12, p=0.0006) and vehicle treated control mice (4.4±9.2%, n=4, p=0.019). Further, there was no difference between the dopaminergic response between LEAP2 treated and LEAP2 treated control mice (p=0.93) or between the two control groups (p=0.99). The area under the curve of the dopamine increase while allowed to eat compared to before eating showed a significant group difference (Figure 4C, One-way ANOVA, F_(3, 15)_=10.07, p=0.0007) where vehicle treated mice had a marked increase (462.5±112.2, n=5) compared to LEAP2 treated animals (67.3±28.5, n=7, p=0.0009) and vehicle controls (56.9±24.1%, n=4, p=0.0025). Notably, LEAP2 treated mice did not significantly differ compared to its control group (52.1±18.3, n=3, p=0.99), nor was there any difference between the two control groups (p>0.99).

**Figure 4.**
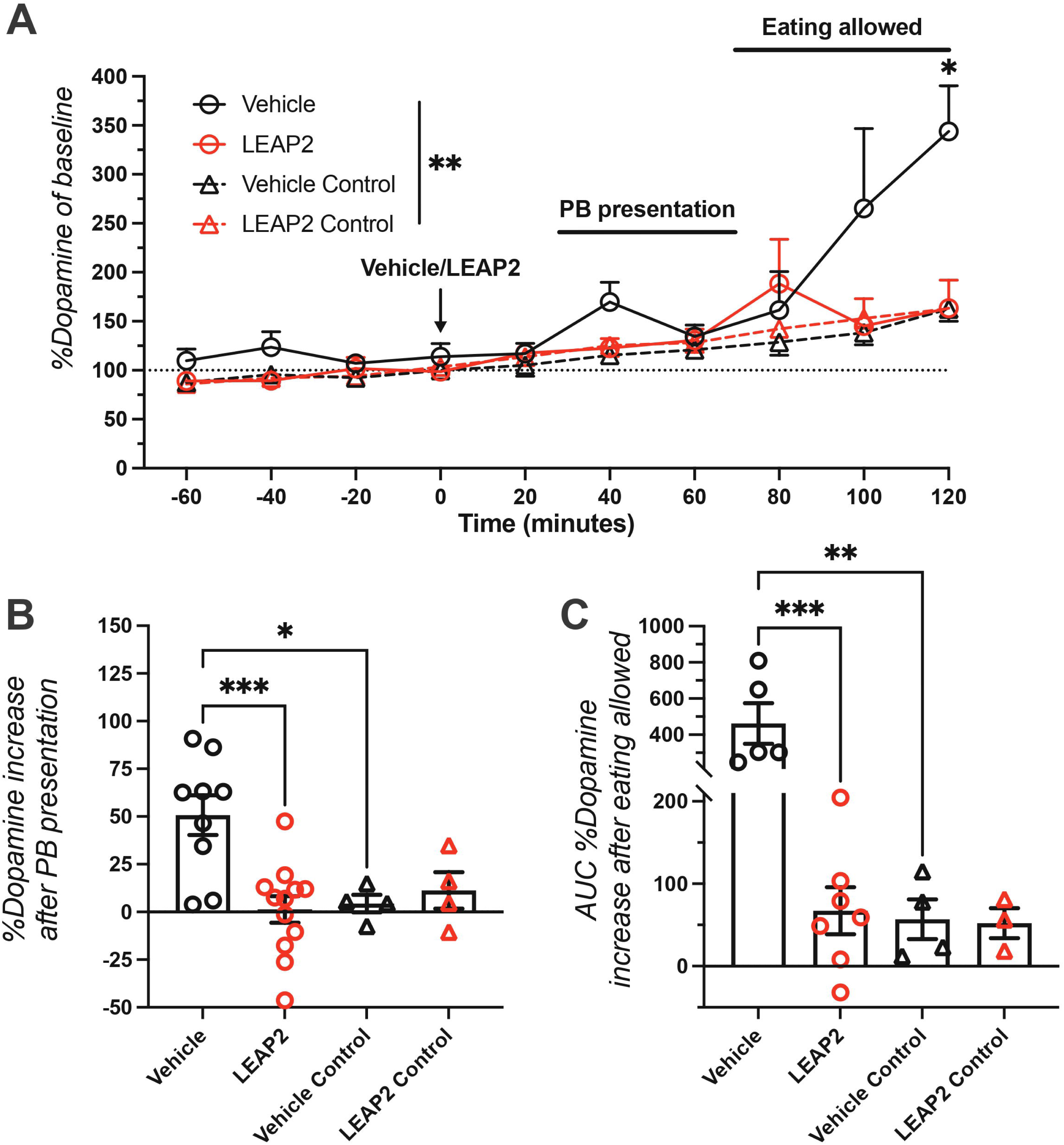
Central LEAP2 abolishes the dopaminergic signaling associated with hedonic feeding. Accumbal dopamine change in response to peanut butter presentation and eating was measured using microdialysis. A group difference (A) between vehicle and LEAP2 treated mice and treated control mice that where not exposed or allowed to eat peanut butter was detected with a significant elevation in vehicle treated mice at 120 minutes. The change in dopamine levels during exposure from just before (B) showed a significant increase only in vehicle treated mice (n=9 and n=4 for vehicle and its corresponding control group, respectively), whereas no difference compared to corresponding control group was detected in LEAP2 treated mice (n=12 and n=4 for vehicle and its corresponding control group, respectively). In a subset of animals that were allowed to eat (C, n=5 and n=7 for vehicle and LEAP2 treated groups, respectively), the dopamine change area under the curve showed a similar pattern where only vehicle treated mice showed a significant increase compared to LEAP2 and control mice. Group differences in A were assessed using mixed-effect analysis with Dunnet’s multiple comparisons test post hoc (p*<0.05 vs. corresponding control and LEAP2 group) and using one-way ANOVA followed by Šidák’s multiple comparisons test post hoc in B and C. Data is shown as mean±SEM and was corrected for multiple comparison. p***<0.001, p**<0.01, p*<0.05 vs. corresponding group.

### Continuous hedonic feeding reduces expression of LEAP2 in reward-related brain areas

Brain areas were punched out and expression of LEAP2 was analyzed from a set of untreated mice to in detail map LEAP2s presence in areas associated with reward and food intake. In chow-fed mice, LEAP2 expression was detected in all investigated brain areas, although at low levels compared to liver and duodenum. One-way ANOVA revealed differences between areas (F_(10, 48)_=48.56, p<0.0001). Notably, the relative expression of LEAP2 was found to be 2.7 and 4.4 times higher compared to hypothalamus in NAc (Table 1, p=0.28) and LDTg (p=0.012), respectively.

**Table 1.** Relative expression of LEAP2 in the brain. LEAP2 expression in different brain areas, liver and duodenum as measured by real-time quantitative PCR. Baseline expression in control mice was compared to expression in hypothalamus as Δ_CT_-values (one-way ANOVA followed by Šidák’s multiple comparisons test post hoc). Mice with free access to peanut butter for one week were compared to the same area in control mice fed a standard chow diet (unpaired t-test).

|  | Baseline expression |  | Peanut butter |  | N |  |
| --- | --- | --- | --- | --- | --- | --- |
|  | vs. hypothalamus |  | vs. chow diet |  |  |  |
| Area | $\Delta_{CT}$ | p-value | $\Delta_{CT}$ | p-value | Chow | Peanut butter |
| PFC | -7.57±0.28 | 0.94 | -8.60±0.48 | 0.093 | 6 | 6 |
| NAc | -6.87±0.65 | 0.28 | -9.23±0.52 | **0.022** | 5 | 5 |
| BNST | -7.70±0.39 | 0.98 | -9.10±0.39 | **0.030** | 6 | 6 |
| Hippocampus | -7.47±0.28 | 0.87 | -9.21±0.39 | **0.0046** | 6 | 5 |
| Amygdala | -7.83±0.47 | 0.99 | -9.78±0.73 | **0.046** | 6 | 5 |
| Hypothalamus | -8.30±0.64 | - | -8.57±0.21 | 0.75 | 6 | 4 |
| VTA | -8.28±0.26 | >0.99 | -9.93±0.22 | **0.00068** | 6 | 6 |
| DR | -7.79±0.36 | 0.99 | -8.20±0.42 | 0.47 | 6 | 4 |
| LDTg | -6.18±0.44 | **0.012** | -9.71±0.31 | **<0.0001** | 6 | 6 |
| Liver | 4.27±0.43 | **<0.0001** | - | - | 3 | - |
| Duodenum | 0.0014±0.96 | **<0.0001** | - | - | 3 | - |

Interestingly, in animals with free access to peanut butter, a strong reduction of LEAP2 mRNA expression could be observed (two-way ANOVA, effect of diet: F_(1, 82)_=59.62, p<0.0001, interaction effect: F_(8, 82)_=2.59, p=0.014), and in particular areas associated with reward, such as NAc (p=0.022), VTA (p=0.00068) and LDTg (p<0.0001), and memory, e.g., hippocampus (p=0.0046). Notably, hypothalamus, which is strongly associated with homeostatic eating and energy homeostasis^3, 33, 34^, had the lowest baseline expression, and was not affected by hedonic feeding (p=0.75)

### Intra-LDTg infusion of LEAP2 attenuates hedonic feeding

The qPCR assay revealed that LDTg had the highest expression of LEAP2 in the brain as well as the greatest effect following prolonged hedonic feeding. Considering its importance for ghrelinergic signaling in reward^8, 9, 12, 35^, LEAP2s effect in LDTg specifically was further investigated. In an acute hedonic feeding paradigm, bilateral infusion of LEAP2 in LDTg successfully reduced peanut butter consumption (Figure 5, repeated measures two-way ANOVA, Treatment effect: F_(1, 11)_=5.63, p=0.037, Time effect: F_(1.00, 11.00)_=17.30, p=0.0016, Interaction: F_(1.00, 11.00)_=0.31, p=0.59). The effect was notable after one hour of peanut butter eating where a significant difference was found between vehicle (221.5±56.6 mg, n=12) and LEAP2 treated animals (119.6±40.7 mg, p=0.0008). However, after two hours of peanut butter consumption this difference was no longer significant (328.0±90.5 mg and 253.7±62.8 mg for vehicle and LEAP2 treated animals, respectively, p=0.42).

**Figure 5.**
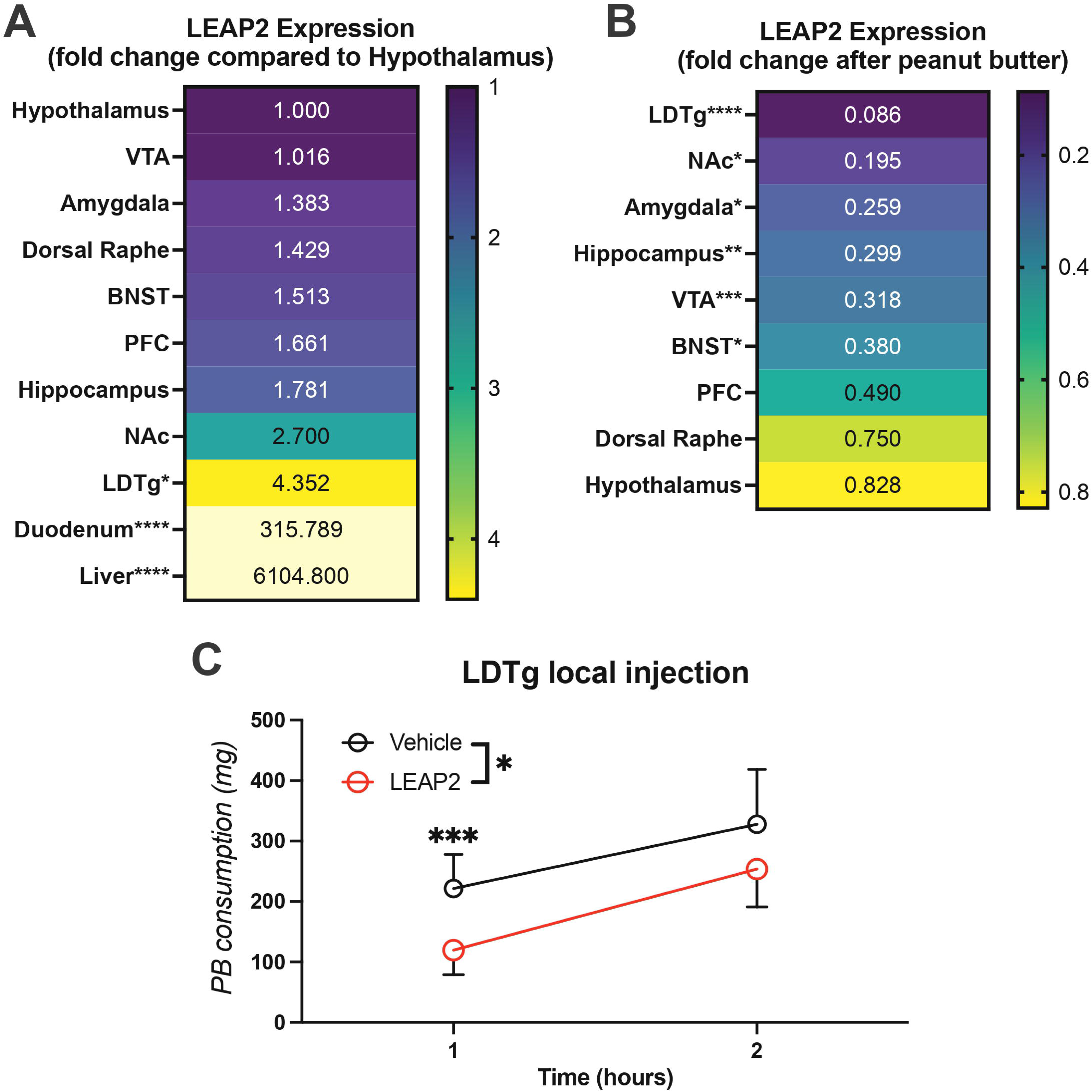
LEAP2 in LDTg is sufficient to reduce hedonic feeding. Out of all analyzed brain areas, LDTg showed the highest expression (A) and was most affected by hedonic feeding (B). Thus, in a binge-eating condition similar to that used in microdialysis and CPP, bilateral infusion of LEAP2 was used to investigate the significance of LDTg for regulating hedonic feeding. For the two-hour duration, a significant group difference could be observed that was significant at the one-hour timepoint (C). However, the effect was not significant two hours after administration. Expression data is shown as fold change and the statistical data is based on the Δ_CT_-values shown in Table 1. For group analysis of hedonic feeding, a repeated-measures two-way ANOVA was used, followed by Wilcoxon test post hoc. Data is shown as mean±SEM and was corrected for multiple comparison. p****>0.0001, p***>0.001, p*>0.05 vs. hypothalamus expression (A) or corresponding control group.

## Discussion

The purpose of the current study was to clarify the role of central LEAP2 on hedonic feeding and how it affects central reward processes in this context. Here, we tested three different palatable food options and concluded not only that LEAP2 reduced hedonic feeding, but it did so in a preference-dependent manner where the mice that showed a higher preference for palatable foods had the strongest effect of LEAP2. Specifically, LEAP2 had the strongest effect on peanut butter consumption, which also had the highest preference. Interestingly, central LEAP2 appears to still affect homeostatic feeding, but secondary to hedonic, since animals receiving chocolate, with a preference close to 50% and no rewarding memory effect, showed a significant reduction in chow intake. Conversely, LEAP2 showed little to no effect on chow intake in peanut butter feeding animals, with a much higher preference. Taken together, this suggests that central LEAP2 reduces feeding generally, but it preferentially affects hedonic rather than homeostatic feeding. This is in line with a recent study showing that central LEAP2 specifically reduces alcohol intake in high-, but now low-consumers^25^. It should be mentioned that although previous studies have shown that central administration of ghrelin increases both chow and hedonic foods^11, 36, 37^ ghrelin increases the incentive value of foods^38^, including low preference foods. Reducing incentive value, e.g., through LEAP2, would thus have little effect on already low preference foods, such as chow when peanut butter is available.

The effect of central LEAP2 to reduce hedonic feeding and food reward was further evident in the CPP experiment. Previously, ghrelin has been shown to increase CPP to high-fat diet in a GHSR-dependent manner^39^. Here, peanut butter caused the strongest CPP, which also showed the greatest reduction after LEAP2 administration, followed by Nutella® and lastly chocolate that provided a minimal CPP with no effect of LEAP2. This paradigm simulated the acute rewarding effect in a better way compared to the *ad libitum* food intake experiment, as they were only allowed to eat for the duration of the conditioning. As such, these results confirm the effect of LEAP2 on *ad libitum* hedonic eating. Interestingly, our expression analysis did reveal a significant reduction of LEAP2 after *ad libitum* peanut butter also in hippocampus along with the several reward-related areas. Taken together, this strongly suggests that LEAP2 have important functions in modulating rewarding memory related to food, an important aspect for food search behaviors which ghrelin is known to affect through hippocampal circuits^40^.

However, it should be noted that due to the experimental setup, we were only able to test rewarding memory retrieval, and not memory formation in a reward CPP setting, which should be considered a limitation in the interpretation of our results.

Further, we investigated how LEAP2 affected dopaminergic reward signaling directly in NAc during peanut butter exposure and eating. In line with previous studies^41^, central LEAP2 did not appear to have any effect on dopaminergic signaling *per se*, suggesting that LEAP2 has no effect on non-rewarding stimuli, e.g., chow and does not affect the normal state. This was further evident in the locomotor activity test where LEAP2 did not affect locomotion. However, LEAP2 successfully attenuated the dopaminergic response to peanut butter exposure as well as peanut butter eating. This is in line with previous studies showing how GHSR antagonists inhibit dopaminergic reward in response to food and different drugs of abuse^8, 13, 42, 43^.

Previous studies have shown that LEAP2 is expressed centrally and can be detected in cerebrospinal fluid^21, 23^. Accordingly, we show that LEAP2 is widely expressed in different brain areas. Strikingly, LEAP2 was highly expressed in areas involved in the reward system and in particular NAc and LDTg. Albeit central LEAP2 expression were at low levels compared to liver and duodenum, it should be noted that while these tissues produce LEAP2 to travel through the bloodstream, the locally produced LEAP2 in the brain may still be relevant to affect the local GHSR in those areas. Indeed, the highest expression of LEAP2 overlap somewhat with where the GHSR is expressed in the brain^14^.

Further, one week of *ad libitum* access to peanut butter greatly reduced the expression of LEAP2, most notably the reward-related areas, such as LDTg and NAc. Seeing how LEAP2 reduced hedonic feeding consequently in this study, a reduction in LEAP2 expression in the brain following *ad libitum* access to peanut butter suggests that central LEAP2 works in a positive feedback loop to enhance hedonic eating. It appears natural that once a high-energy and rewarding food has been found, the brain would act to enhance that sensation and memory to motivate the search and consumption of it^44, 45^. However, in modern humans this creates a vicious eating cycle of high- and fast-energy food with high fat and/or sugar content, leading to weight gain^46^.

Considering LEAP2s high expression in reward-related areas and the effect caused by a short period of hedonic feeding, as well as the profound acute effect on hedonic feeding and reward, it seems clear that central LEAP2 is an important modulator of dopaminergic reward signaling. Exactly how LEAP2 reduces dopaminergic reward signaling is uncertain, but likely involves, at least in part, a reduction of cholinergic input to VTA from LDTg, since local injection of LEAP2 here was associated with reduced hedonic feeding. Indeed, previous studies have shown ghrelin concomitantly induces accumbal dopamine release and acetylcholine release in VTA, an effect that can be blocked by GHSR antagonists^9^. Further, ghrelin-induced feeding and accumbal dopamine release can be attenuated by blocking cholinergic signaling^12, 47^.

Although not tested here, it appears also credible that LEAP2 may act to reduce dopaminergic signaling directly through the GHSR-D2 receptor complex, which could reduce presynaptic and somatodendritic dopamine release in NAc and VTA^24^. In particular since LEAP2 showed high expression and peanut butter effect in NAc and VTA. Surprisingly, in this study, hypothalamic LEAP2 appeared to be the least important for hedonic feeding, considering its low LEAP2 expression and that free hedonic feeding did not alter that expression. A previous study has shown that overexpression of LEAP2 in hypothalamus reduces both chow and high-fat diet consumption^34^, an effect thought to be mediated through arcuate nucleus proopiomelanocortin neurons. Therefore, it appears likely that, although not found here, hypothalamic LEAP2 would affect food intake. This discrepancy might be explained by the difference between hedonic and homeostatic feeding. Here, the focus was on the rewarding aspects of hedonic feeding and these behaviors are reliant mainly on central reward signaling, whereas hypothalamic GHSR modulation is instead mainly associated with the homeostatic aspect of food intake^3, 33^. As such, our results further strengthen that central LEAP2 preferentially affects the rewarding component of eating, at least when a hedonic palatable food option is available.

Beyond feeding, the finding that central LEAP2 can attenuate dopaminergic reward signaling warrants further studies of its effect in different dopamine-related psychiatric disorders, most distinctively addiction disorders where GHSR signaling already have been shown to be affected^17–19, 48^. Interestingly, LEAP2 appears greatly affected by diet, as shown here and in previous studies^49, 50^, as well as by the gut microbiome^51^. Importantly, we show here that even LEAP2 expression in deep brain structures are highly affected by diet. Although the exact nature of dietary and microbiota regulation of LEAP2 remains to be fully elucidated, it opens several avenues by which LEAP2 levels can be manipulated and as such, several options for treating dopamine-related psychiatric disorders through LEAP2. For instance, a recent study showed that central LEAP2 reduces alcohol intake^25^.

However, for the translational value of the present study, a couple of limitation exists. As such, the current study was limited to only using male mice. Considering that there appears to be some sex-dependent differences in LEAP2^51, 52^, any such differences must be addressed to fully understand the effects of LEAP2. Further, we aimed to investigate the effects of acute central effects of LEAP2, and not the effects of systemic or prolonged elevated LEAP2, which further studies should address.

In conclusion, we here show that central LEAP2 attenuates the dopaminergic reward and reward-related behavior that is associated with hedonic eating. This effect appears mainly mediated through LEAP2 within the dopaminergic reward pathway, including cholinergic modulation from LDTg.

## Acknowledgement

This study was supported by grants from the Swedish Research Council (2019-01676), LUA/ALF (grant no. 723941) from the Sahlgrenska University Hospital and the Swedish brain foundation (EJ and MT-A (PS2022-0026)).

Ellen Hjertkvist did not meet all criteria for authorship, but is gratefully acknowledged for valuable technical assistance.

