## Supplement figures for "Decoding the Influence of Central LEAP2 on Hedonic Food Intake and its association with Dopaminergic Reward Pathways"

**Supplementary Figure 1. Injection and probe placements.**

Schematic representation of the target for injection and probe placements, stereotaxic coordinates, and examples of placements. A) Intracerebroventricular injections, B) bilateral injections in the LDTg and C) probe placement in the nucleus accumbens shell.

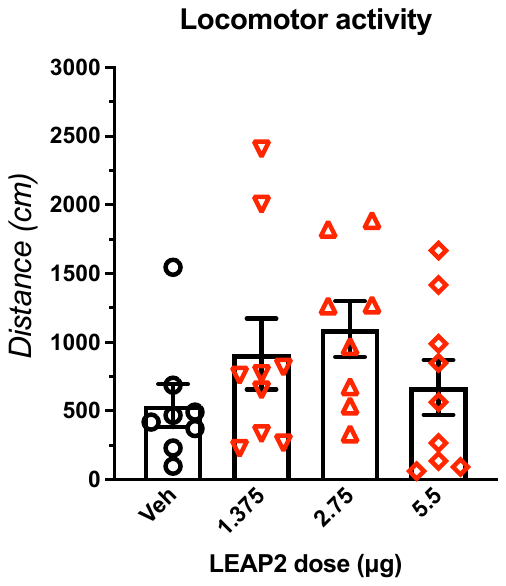

**Supplementary Figure 2. Dose-response on locomotor activity.**

The effect of centrally administered LEAP2 was assessed at doses of 1.375, 2.75 and 5.5 μg on locomotor activity. The dose of 5.5 μg was deemed to have the least effect on the general state of the animal. Group comparisons were made using one-way ANOVA. Data is shown as mean±SEM. No significant difference was found (F_(3, 30)_=1.34, p=0.28).
